# Beyond the Barrier: An Assessment of Social Behaviour in Mice using a Modified 3-Chamber Test

**DOI:** 10.64898/2026.01.30.702763

**Authors:** Meida Sofyana, Hugh D. Piggins, Megan G. Jackson, Emma S.J. Robinson

## Abstract

**Background:** The three-chamber test (3CT) is widely used to assess social behaviour in mice, based on the assumption that time spent near a conspecific reflects motivation for social contact. However, the design of the task constrains interpretation, as behaviour may reflect exploration, novelty seeking, or territorial investigation rather than affiliative social motivation. In addition, key biological factors such as sex differences and social hierarchy are often overlooked.

**Aims:** We hypothesised that the 3CT overestimates sociability and used a direct-interaction phase to investigate motivation for affiliative social contact. We also integrated social status to determine if this modulated behavioural patterns and interacted with sex.

**Methods:** Adult male and female C57BL/6 mice (n = 32) were tested in a standard 3CT, followed by removal of the cage barrier to permit direct contact. Behavioural parameters were quantified, and social status was determined using the tube test.

**Results:** Males exhibited higher social interest index scores than females. Once the barrier was removed, both sexes displayed a negative direct sociability index, indicating greater environmental exploration than social engagement. Correlation analysis revealed no association between indirect and direct measures. Sex differences emerged primarily among submissive mice, with submissive males showing greater social investigation than submissive females.

**Conclusion:** These findings suggest that standard 3CT indices reflect exploratory rather than affiliative social motivation. The modified paradigm incorporating direct interaction provides a more realistic assessment of social behaviour and challenges assumptions about intrinsic sociability in mice.

## 1. Introduction

The three-chamber test (3CT) is widely used to assess social behaviour in mice (Rein & Yan, 2020). In this test, a mouse is placed in an arena with three interconnected compartments. The mouse can freely explore, and its social behaviour is assessed by observing its preference to spend time in the chamber with a social stimulus (familiar or novel, adult or juvenile mouse) rather than with a non-social stimulus (an object or an empty chamber). Typically, these stimuli are placed inside cages to minimise physical interaction (Yang et al., 2011; Crawley, 2004). The 3CT can be used as a single test or repeated to assess social memory (Leung et al., 2018; Kaidanovich-Beilin et al., 2011).

In the simplest version of the task (object/empty chamber versus unfamiliar mouse), exploration of the social stimulus is used to define sociability, with greater exploration of the cage containing the mouse interpreted as higher social interest (Fairless et al., 2011). Mice are widely regarded as social animals that rely on frequent investigation and contact to maintain social relationships (de la Zerda et al., 2022). Their interactions are strongly guided by olfactory cues, which facilitate individual recognition, dominance assessment, and mate selection (Muna et al., 2023; Thompson et al., 2018).

Recent studies challenge the assumed sociability of mice, particularly in relation to male mice. Renaturalisation studies in mice suggest that, when provided with sufficient space and resources, social interactions are reduced, particularly in males (Vogt et al., 2024). Tallent et al. (2024) also found that male mice housed in partially divided cages displayed lower anxiety and reduced aggression compared to those in standard cages, and a recent study quantifying affective state in single versus group-housed male mice found that those housed individually were in a more positive affective state (Davies et al., 2025).

Interaction with a social stimulus in the 3CT is widely interpreted as motivating positive social contact (Arakawa, 2023). However, exploration, social dominance, and aggression associated with territorial behaviour could also drive social interactions (Shemesh et al., 2024). Social hierarchies play an important role in mouse social behaviour, with dominant and submissive individuals exhibiting distinct patterns of social engagement (Fetcho et al., 2023). Dominant mice may assert their status through increased interaction, while submissive mice may engage less or exhibit avoidance behaviours (Kunkel & Wang, 2018). Therefore, aggression and the social hierarchy of test and stimulus mice could also lead to differences in behaviour in the 3CT. Studies using the 3CT rarely account for hierarchical differences, and ignoring mice’s dominance status may obscure important behavioural nuances (Sapolsky, 2005).

In this study, we first conducted a systematic review of 3CT protocols used to assess social behaviour in male and female mice in order to identify methodological heterogeneity and determine the most commonly applied experimental parameters. Although the general structure of the 3CT was broadly consistent across studies, substantial variation was evident in habituation procedures, stimulus characteristics, and behavioural outcome measures, all of which may influence interpretation and reproducibility (Jabarin et al., 2022; see Supplementary Figs. 1 and 2). Based on these findings, we implemented a standard 3CT protocol using the most frequently reported methodological parameters to ensure comparability with the existing literature.

However, because conventional 3CT measures infer sociability from interaction with a physically restricted stimulus, we questioned whether proximity-or cage-directed behaviours truly reflect motivation for positive social contact. To directly test this assumption, we modified the protocol by lifting the cage barrier at the end of the session to permit unrestricted interaction between the test mouse and the social stimulus. This approach allowed us to (a) evaluate whether exploration of a confined conspecific predicts affiliative motivation once direct contact becomes possible, and (b) determine how behavioural priorities change when the social stimulus is fully accessible. In addition, we incorporated social hierarchy into our analysis by assessing dominance status using the tube test, enabling us to examine how social status modulates social behaviour and interpretation of standard 3CT metrics.

## 2. Methods

### 2.1. Systematic Review of 3CT Protocols

The systematic review focused exclusively on studies involving mice that employed the three-chamber test to assess social behaviour. To minimise duplication and reduce reporting bias, the review protocol was prospectively registered on PROSPERO (CRD42024557108) (Sofyana et al., 2024). PubMed and Embase databases were searched from inception to July 2024 using the terms “three-chamber test,” “3-chamber test,” and “three-chambered test.” Eligible studies included original research articles, reviews, and systematic reviews published in English with full-text availability to enable comprehensive data extraction and accurate interpretation.

Screening and prioritisation were guided by relevance to the experimental design, with the highest priority given to animal studies using rodents that reported methodological parameters of the three-chamber test. Publication characteristics (e.g., peer-review status and publication date) were considered secondary. Data extracted from the review were used to identify the most frequently applied protocol features and behavioural outcome measures, which informed the design of the standard three-chamber protocol implemented in the present study (see Supplementary Figures 1–2 and Sofyana et al., 2024 for full methodological details and results).

### 2.2. Social Behavioural Analysis

#### 2.2.1. Animals

A sample size of 16 mice per sex was determined based on an a priori power analysis using G*Power 3.1.9.7. Expected effect sizes were in the moderate-to-large range (Cohen’s f = 0.35–0.7), with an α error probability of 0.05 and statistical power set at 0.8 (1–β). This analysis indicated that a total sample size of 32 animals was appropriate to detect a main effect of sex and social hierarchy (Szabó et al., 2024; Fairless et al., 2011). Male and female C57BL/6JOlaHsd mice, 5 weeks of age upon arrival, were purchased from Envigo RMS Limited (Bicester, Oxfordshire, United Kingdom). Mice were pair-housed with enrichment consisting of nesting material (50% paper bedding (IPS, UK) and 50% Sizzlenest bedding (Datesand, UK), one cardboard tube suspended from the cage lid, two cardboard tubes, a wooden ball, and a chew block. They were kept in temperature-controlled conditions (20–21 °C, 45–55% relative humidity, 12:12-hour light-dark cycle) under a 12:12-hour light-dark cycle (lights-on at 08:15 and lights-off at 20:15). Standard lab chow (Purina, UK) and water were provided ad libitum. Weights were monitored weekly. Mice spent three weeks acclimatising to the facility before the start of the experiment, and we habituated them to handling by one female experimenter (Sorge et al., 2014) and managed them in accordance with the 3Hs protocols (3Hs Initiative, 2024). Mice were 8 weeks old at the start of the experiment (male: 24.2–26.3 g; female: 19.0–21.6 g), consistent with the predicted age when stable social status develops in the C57BL/6 strain (Battivelli et al., 2024). Experiments were performed in accordance with the Animals (Scientific Procedures) Act (UK) 1986 and were approved by the University of Bristol Animal Welfare and Ethical Review Body (AWERB).

#### 2.2.2. The Three-Chamber Apparatus

The test apparatus comprised a rectangular transparent Plexiglas box (L60 x W40 x H22 cm) divided into three equal chambers (see Supplementary Figs. 2C and 2D). Opaque black acrylic dividing walls had small square doors (L7 x W5 cm) that provided unrestricted access to all chambers. The entire arena was set on a white Plexiglas floor to provide a contrasting background for video recording. The central chamber was empty, and each side chamber contained a round metal-barred cage (a diameter of 10 cm and a height of 10 cm) with 0.4 mm bars spaced 1 cm apart (Contestabile et al., 2021), allowing visual and olfactory contact but preventing direct physical interactions (Jiang et al., 2015; Bagosi et al., 2017). A lid was secured on top of the cage to prevent the test mouse from gaining entry. The cage was placed in one-half of the chamber, opposite the door, and there was sufficient space between the cage and the apparatus’s outer wall, allowing the test mouse to explore the cage’s perimeter fully (Figure 1).

**Fig.1.**
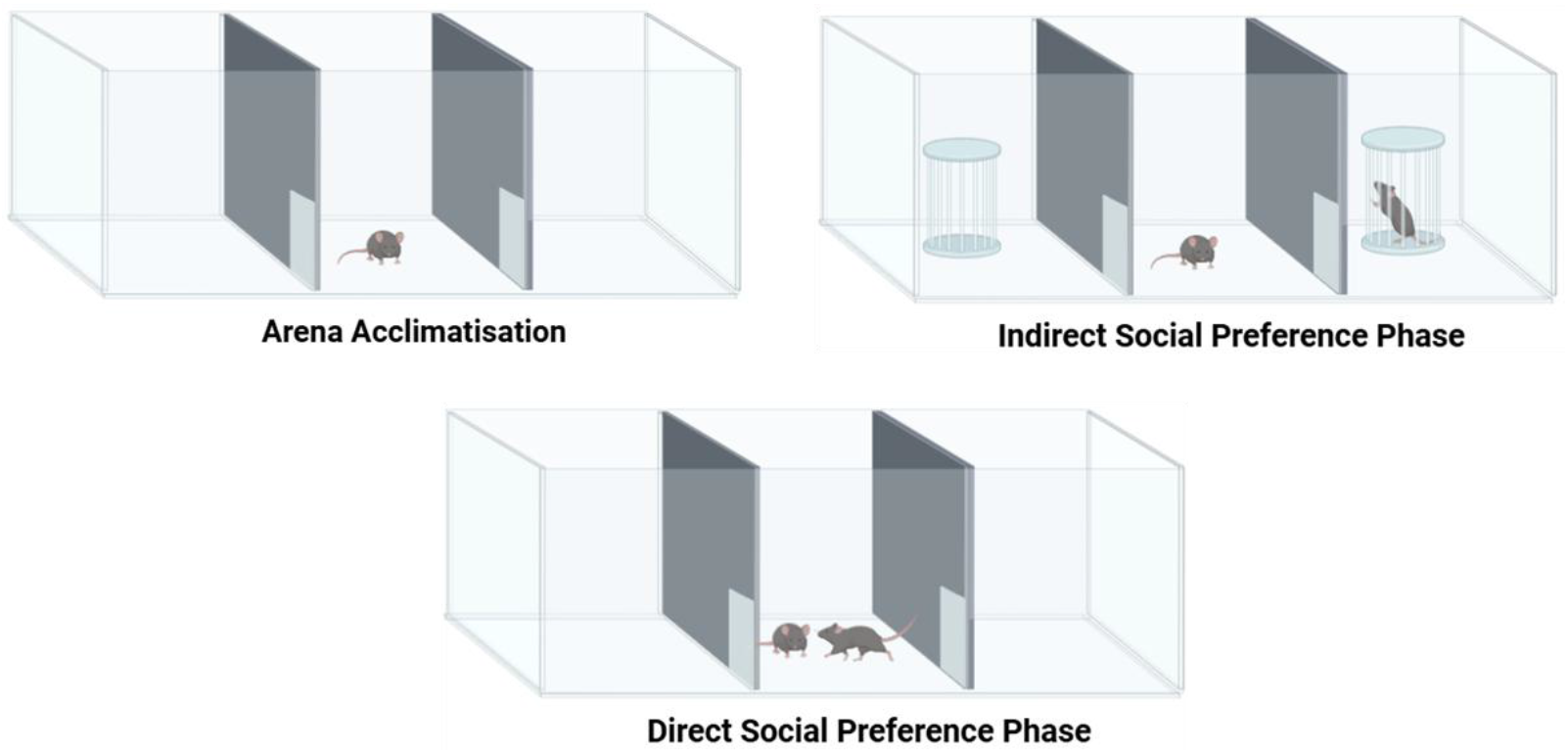
The Modified Three-Chamber Test. The test consists of 10 minutes of arena acclimatisation, 7 minutes of the indirect interaction phase, and 3 minutes of the direct interaction phase. During this direct interaction phase, both the test and stimulus mice were allowed to explore the entire chamber. (Created with BioRender.com)

#### 2.2.3. The Modified Three-Chamber Test

The protocol used in this experiment followed the most widely used method, as identified in a review of 264 studies. All experiments were conducted during the animals’ inactive phase in a quiet room (180 lux; see Supplementary Figs. 2I and 2J), measured with the Lux Light Meter Pro App v1.8.5 (webrain.ltd). All chambers were evenly illuminated, ensuring no shadows (Kurahashi et al., 2024). The apparatus’s location and orientation within the test room remained fixed. Testing was conducted in two independent batches (n = 16 per batch), each consisting of 8 males and 8 females. To minimise order effects, the sequence in which males and females were tested was counterbalanced across days, ensuring that neither sex was consistently tested earlier or later in the session. The position of the social-stimulus mouse (left vs. right chamber) was also counterbalanced across subjects to prevent the development of a side bias.

The mice were acclimatised to the arena immediately preceding the test (see Supplementary Fig. 2F). The test mice were placed in the middle chamber and allowed to explore the entire empty chamber for 10 minutes (see Supplementary Figs. 2G and 2H). Animals that spent more than 80% of the 10-minute acclimatisation phase exploring either side chamber were excluded from data analysis (Piccin et al., 2022). No side bias was detected during acclimatisation in our experiment. Immediately following acclimatisation, an unfamiliar same-sex stimulus mouse was placed in a cage in one side chamber (social chamber), and an empty cage was placed in the other (non-social chamber; see Supplementary Fig. 2K). The test mouse was allowed to explore the arena and cages for 7 minutes to assess social preference for the social or non-social stimulus (see Supplementary Fig. 2L). After 7 minutes of indirect contact, both cages were lifted, and behaviour was observed for a further 3 minutes (Figure 1).

The stimulus mouse was unfamiliar but strain-, sex-, weight-, and age-matched with the tested mice. The stimulus mice had never been in physical contact with the subject mice (Gao et al., 2020), and from a cage which had not had direct visual contact (Ferreira, 2025; Greer et al., 2025; Ueno et al., 2024). During the test, the starting point for the stimulus mouse was counterbalanced across tests to account for potential innate place preference (IPP) and to reduce odour accumulation. The entire test period was recorded by a video camera (Logitech C920 HD Pro Webcam) placed 150 cm above the apparatus. The videos were analysed offline and scored manually (Zahran et al., 2024) using Novel Object Video Counter (https://novel-object-video-counter.netlify.app/). The experimenter was blind to social status (dominant or submissive) during behavioural scoring. The floor and walls were cleaned between each test with water and 70% ethanol, and dried with a paper towel to reduce olfactory cues. To reduce the transfer of novel social odours acquired during the test session, tested mice were kept separate from their untested cage mates in a temporary cage until all animals from the same home cage had been tested (Drapeau et al., 2018; Lin & Hsueh, 2014).

Details of the recorded behaviours are presented in Table 1, and CSI and related metrics are calculated as shown in Figure 2. A sociability index of zero indicates no preference between the social and non-social cues. A positive index reflects a preference for the social cue, while a negative index suggests a preference for the non-social cue.

**Figure 2.**
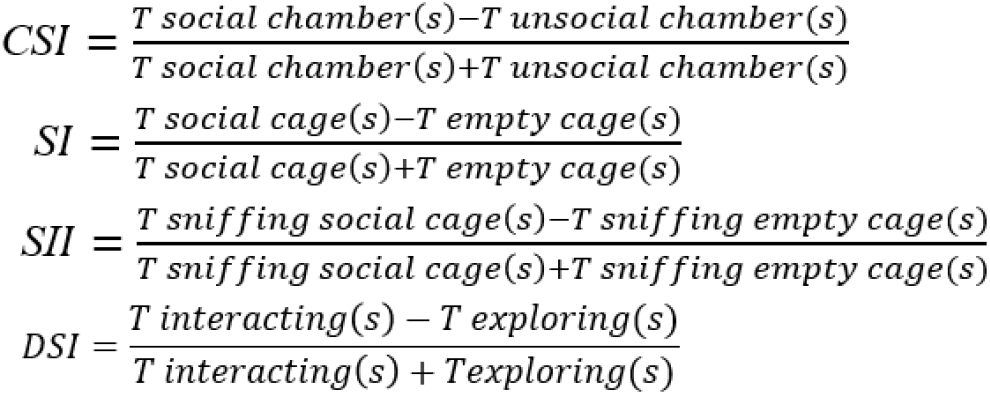
Definitions of sociability indices used in the three-chamber test. The Chamber Sociability Index (CSI) represents the relative time spent in the social versus non-social chamber. The Sociability Index (SI) reflects the proportion of time spent near the social cage relative to the empty cage. The Social Interest Index (SII) quantifies the relative time spent sniffing the social cage compared with the empty cage. The Direct Sociability Index (DSI) represents the relative time spent directly interacting with the stimulus mouse versus environmental exploration during the direct contact phase. All indices range from -1 to +1, with positive values indicating a preference for the social stimulus and negative values indicating a preference for the non-social stimulus. T indicates total time in seconds.

**Table 1.** The variables observed during the first 7 minutes. **Table 1. Behavioural variables quantified during the indirect phase of the three-chamber test (first 7 min).** The table summarises the parameters extracted during the indirect (cage-based) phase of the three-chamber test, including measures of general exploration, investigatory behaviour, and commonly used sociability indices.

| Parameter | Abbreviation | Definition / Description | Behaviour Assessed |
| --- | --- | --- | --- |
| Total transitions between chambers | NA | Number of times the mouse moves between the social and non-social chambers | General activity/exploration |
| Frequency near each cage | NA | Number of times the mouse is within proximity of each cage | Interest/approach behaviour |
| Frequency sniffing each cage | NA | Number of times the mouse sniffs each cage | Social investigation |
| Latency to enter each chamber | NA | Time taken to first enter a social or non-social chamber | Exploration behaviour |
| Latency to approach the cage | NA | Time taken to approach the cage | The initiation of approach behaviour |
| Latency to sniff the cage | NA | Time taken to start sniffing the cage | The initiation of investigation behaviour |
| Chamber Sociability Index | CSI | Relative time spent exploring the social chamber versus the non-social chamber | Social preference |
| Sociability Index | SI | Proportion of time the mouse's nose is within 3 cm of the social cage | Social preference |
| Social Interest Index | SII | Relative time spent sniffing the social cage | Social investigation |
| Direct Sociability Index | DSI | Relative time spent interacting directly with the social stimulus | Social interaction |

The parameters observed during the last 3 minutes are presented in Table 2, that were scored and categorised as: social investigation behaviour (all forms of olfactory investigation, where mice sniff to gather social and chemical information about the stimulus mice), social interaction and contact behaviour (involve direct physical contact or proximity between mice that demonstrate active interaction, engagement, and movement towards or around another mouse), exploratory behaviour (behaviours are generally directed towards the environment rather than stimulus mouse, including wall climbing and rearing; rising on hind legs sniffing the air), self-maintenance behaviour (behaviours are generally directed towards the self rather than stimulus mouse, like self-grooming, cleaning fur or paw scratches), defensive behaviour (non-affiliative interactions that are potentially harmful or stress-inducing, like freezing and direct conflict behaviour or active fighting such as punching or biting). Only behaviours from the test animal were used in the analysis.

**Table 2.** The variables observed during the last 3 minutes. **Table 2. Behavioural ethogram used to quantify direct social and non-social behaviours during the final 3 min of the modified three-chamber test.** Following removal of the barrier, behaviours were classified into investigatory, interaction, exploratory, self-directed, and defensive categories. Scoring was restricted to behaviours performed by the test mouse to isolate its behavioural response to unrestricted social access.

| **Behaviour Category** | **Definition / Description** |
| --- | --- |
| Social Investigation Behaviour | All forms of olfactory investigation, where the mouse sniffs to gather social and chemical information about the stimulus mouse |
| Social Interaction and Contact Behaviour | Involve direct physical contact or proximity between mice that demonstrate active interaction, engagement, and movement towards or around another mouse |
| Exploratory Behaviour | Behaviours generally directed towards the environment rather than the stimulus mouse, including wall climbing and rearing (rising on hind legs, sniffing the air) |
| Self-Maintenance Behaviour | Behaviours generally directed towards the self rather than the stimulus mouse, e.g., self-grooming, cleaning fur, paw scratches |
| Defensive Behaviour | Non-affiliative interactions that are potentially harmful or stress-inducing, like freezing, direct conflict, or active fighting (punching, biting) |

Note: Only behaviours from the test animal were tabulated and used in the analysis

#### 2.2.4. The Tube Test

We conducted the tube test at 13 weeks of age, after 8 weeks of cohabitation. We used a 20-cm-long tube for habituation and a 30-cm-long tube for the actual tube test. The tube diameter was adjusted according to sex (2.5 cm for females and 3.5 cm for males) to account for differences in body size and to prevent mice from passing each other, thereby ensuring that outcomes reflected pushing behaviour rather than avoidance or overtaking (Figure 3).

**Fig.3.**
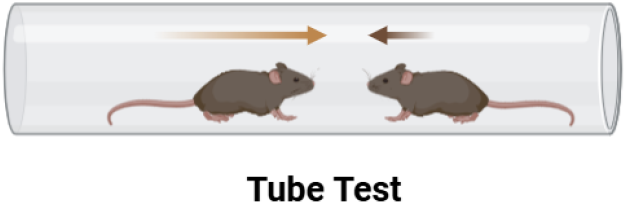
Tube test to assess the social status. In this test, two mice enter the tube from opposite ends. The mouse that can push its opponent out of the tube is considered the dominant individual. (Created with BioRender.com)

The test followed a three-phase protocol: (1) The mice were habituated with a 20-cm tube for 1 hour per day in the home cage for 3 days. Mice were also acclimated to the testing environment to reduce stress and ensure natural behaviour during subsequent phases, (2) mice were guided through the tube 10 times daily for 2 days, five times from each side. (3) mice were tested for 4 days, 4 trials per day, when pairs of mice from the same cage were placed at opposite ends of the tube and allowed to interact. The dominant mouse is identified by its ability to push the submissive mouse out of the tube on at least 3 out of 4 trials and over 3 out of 4 days.

### 2.3. Statistical Analysis

Data normality was tested using the Shapiro–Wilk test. Where data were normally distributed, Repeated Measures (RM) two-way ANOVA, one-way ANOVA, or an independent t-test was performed where appropriate. Normal data are presented graphically as the mean ± standard error of the mean (SEM). When data were not normally distributed or required a two-factor RM analysis, the Mann–Whitney U or Kruskal–Wallis tests were used, and data were presented as median and interquartile range. Significant main effects were defined as p < 0.05 and were followed up with appropriate post hoc analyses, which were then followed by multiple comparisons. A Holm-Šídák pairwise comparison was used following RM two-way ANOVA analysis, Tukey’s post hoc comparison was used following one-way ANOVA analysis, and Dunn’s post hoc comparison was used following the Kruskal–Wallis test. To assess the relationship between indirect and direct social interaction scores, correlation analyses were performed using Pearson’s or Spearman’s rank correlation, depending on the normality of the data. Correlation coefficients (r) and corresponding p-values were reported for each group. Statistical analysis and graphs were created using GraphPad Prism v10.2.0 (Boston, Massachusetts, USA).

## 3. Results

### 3.1. Sex Differences in Exploratory Activity and Social Preference Measures

The absence of an innate side bias was confirmed during the initial 10-minute acclimatisation phase in the empty arena (Supplementary Fig. 3). During the acclimatisation phase, female mice exhibited significantly more transitions between chambers than males (t_(30)_= 3.116, p = 0.004; Fig. 4A).

**Fig.4.**
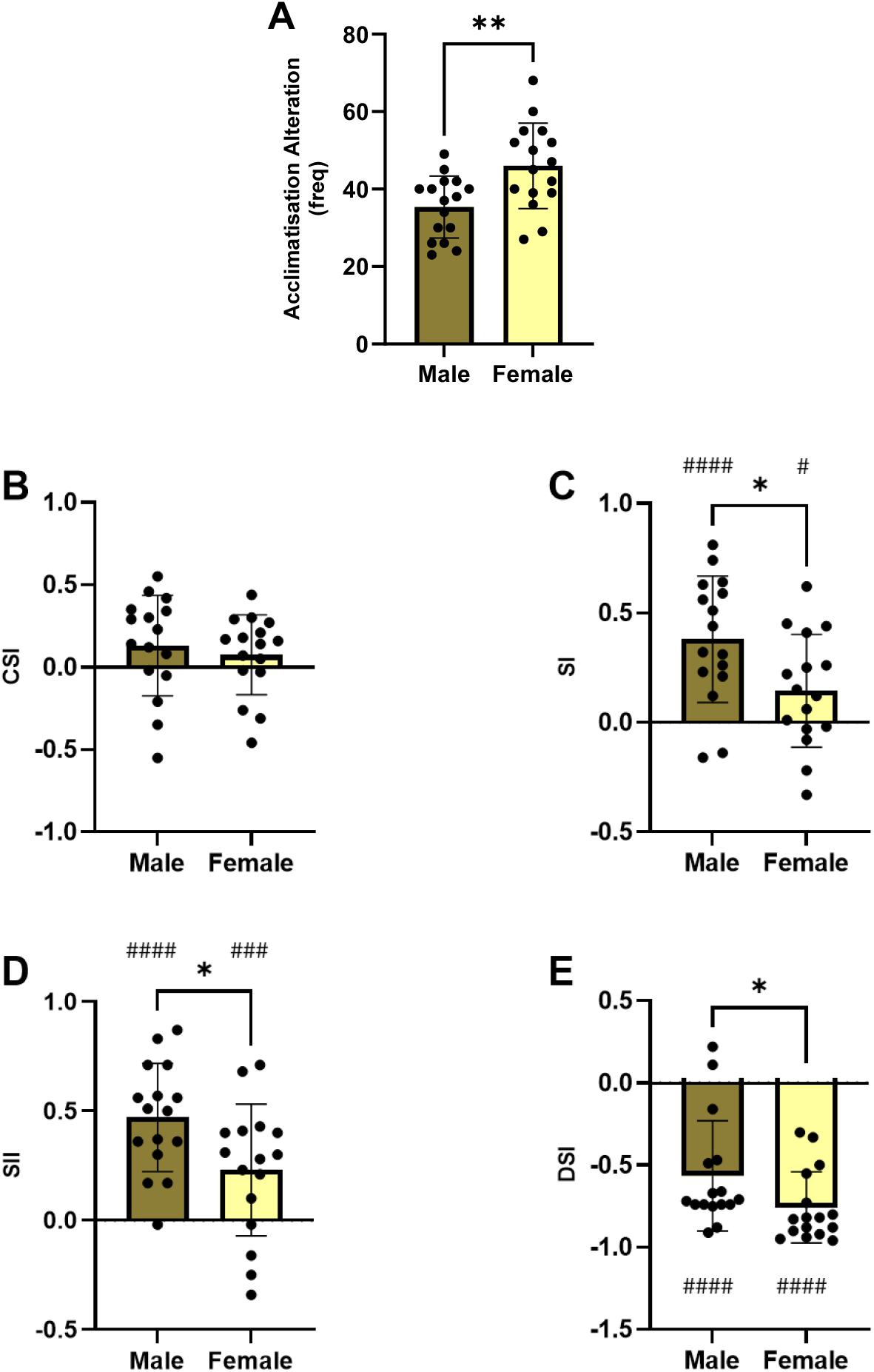
Males have a higher social motivation and strong social interest, and females exhibit higher exploration in the 3CT. **A** Female showed more alteration between social and unsocial chambers during the acclimatisation phase **B** There was no difference in the chamber sociability index between sex groups (p>0.05, independent t-test) **C** Males spent more time near the social cage, and **D** males spent more time sniffing the social cage. Both sexes spent more time exploring, but males spent more time interacting with social stimuli compared to females (#p<0.05, ###p<0.001, ####p<0.0001, one-sample t-test; *p<0.05, **p<0.01, independent t-test).

Neither male nor female mice showed a significant CSI (one sample t-test p>0.05) and there were no sex difference was observed in the chamber sociability index (CSI; p > 0.05; Fig. 4B) however, males showed a higher SI than females (t_(30)_= 2.430, p = 0.0213; Fig. 4C). One-sample t-tests confirmed that SI values were significantly greater than zero in both males (t_(15)_= 5.266, p < 0.0001) and females (t_(15)_= 2.238, p = 0.0409), indicating an overall preference for proximity to the social stimulus. Similarly, males displayed a higher SII than females (t_(30)_= 2.462, p = 0.0198; Fig. 4D), and SII values were significantly greater than zero in both sexes (males: t_(15)_= 7.615, p < 0.0001; females: t_(15)_= 3.059, p = 0.008).

Following removal of the barrier, both sexes exhibited a negative DSI, indicating a behavioural shift toward environmental exploration rather than interaction with the freely moving stimulus animal. One-sample t-tests confirmed that DSI values were significantly below zero for both males (t_(15)_= 6.739, p < 0.0001) and females (t_(15)_= 13.96, p < 0.0001). A Mann–Whitney test revealed a significant sex difference, with females exhibiting a lower DSI than males (median DSI: males = –0.715, females = –0.825; U = 67, p = 0.0203; Fig. 4E), indicating higher levels of non-social exploration in females following barrier removal.

### 3.2. No Sex Differences in Latency or Frequency Measures during Indirect and Direct Social Phases

Analysis of latency and frequency measures revealed no significant sex differences in basic social approach behaviours during either the indirect or direct phases of the task. During the indirect phase, males and females did not differ in latency to enter the social chamber (Supplementary Fig. 4A) or latency to approach the social stimulus cage (Supplementary Fig. 4B). Similarly, during the direct interaction phase, there was no sex difference in latency to approach the social stimulus following barrier removal (Supplementary Fig. 4C).

Consistent with these findings, no sex differences were observed in frequency-based measures of social approach. Males and females showed comparable numbers of entries into the social chamber (Supplementary Fig. 5A), visits to the social cage (Supplementary Fig. 5B), and cage-directed sniffing events (Supplementary Fig. 5C) during the indirect phase. Likewise, the frequency of direct interaction events during the direct contact phase did not differ between sexes (Supplementary Fig. 5D).

### 3.3. Male Mice Exhibit More Social Investigation and Contact Behaviours During Direct Interaction

During the direct interaction phase, based on the median of total activity, males accounted for 34% of total activity in social investigation behaviours and 10% in social interaction/contact behaviours. In contrast, females allocated 19% and 4% to these categories, respectively. Females devoted a greater proportion of activity to exploratory behaviours (71%) than males (52%). Activity in self-maintenance (males 2%, females 0%) and defensive behaviours (males 2%, females 6%) was minimal in both sexes (Fig. 5A).

**Fig.5.**
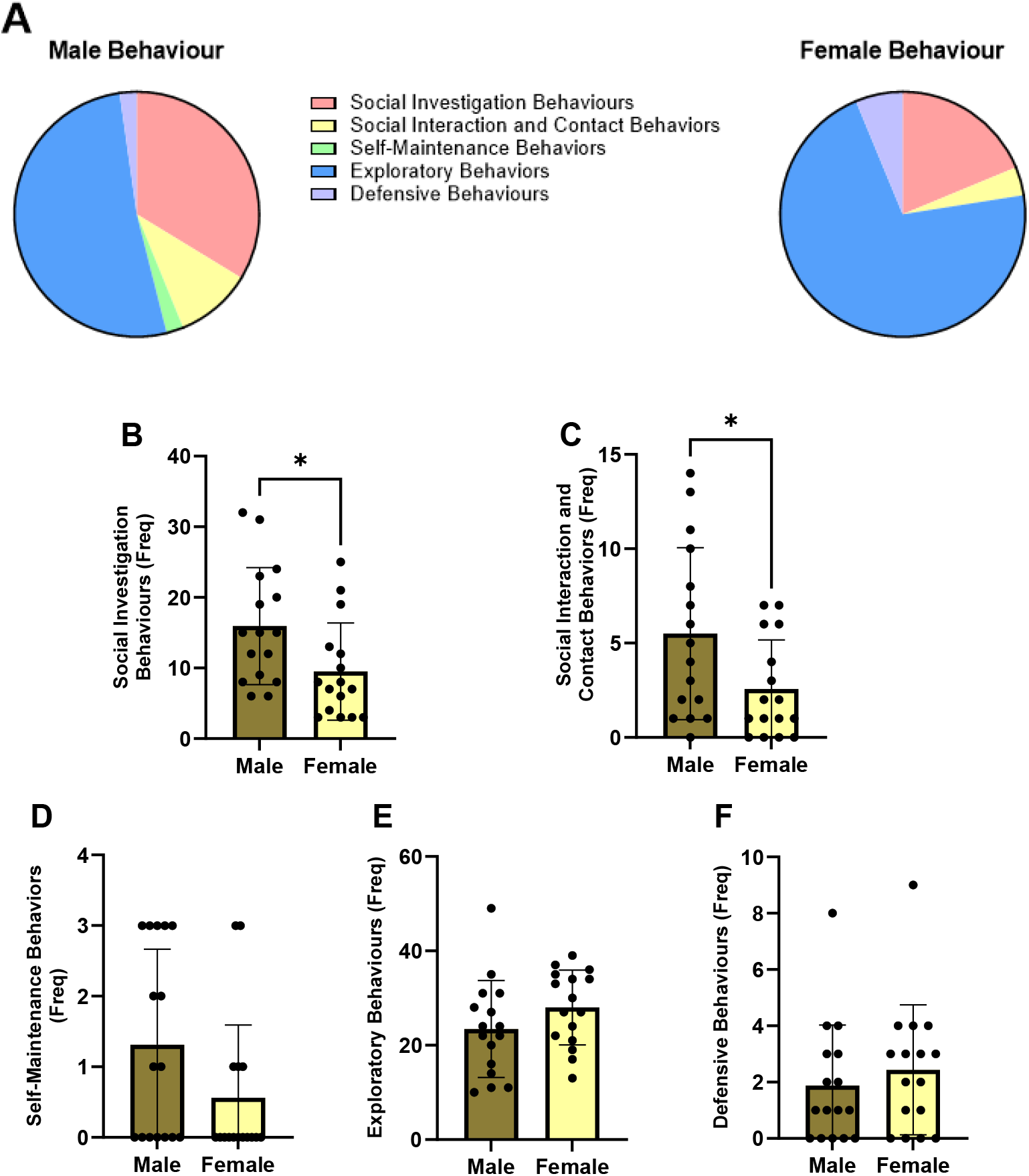
Behaviour of male versus female mice during the three-chamber test without the cage. **A** The quantification of the behaviour shown by males and females during direct social interaction with the stimulus mouse. Male groups engaged more in **B** social investigation and **C** contact behaviours, but no difference between males and females in **D** self-maintenance behaviour, **E** exploratory behaviours, and **F** defensive behaviours (*p<0.05, independent t-test).

Quantitative analysis showed that males spent significantly more time engaging in social investigation behaviours than females (t_(30)_= 2.390, p = 0.0233; Fig. 5B) and also exhibited greater social interaction/contact behaviour (male median = 4.5, female median = 1.5; U = 76, p = 0.048; Fig. 5C). No sex differences were observed (p > 0.05) for self-maintenance behaviours (Fig. 5D), exploratory behaviour (Fig. 5E), or defensive behaviour (Fig. 5F).

### 3.4. Sex Differences in Behaviour Emerge Among Submissive but Not Dominant Mice

When examining the effects of sex and social status on chamber crossings during the acclimatisation phase, two-way ANOVA revealed a significant main effect of sex (F (1, 28) = 10.02, p = 0.0037), but no significant main effect of social status and no sex × social interaction (p > 0.05). Post hoc comparisons indicated that the overall sex effect was driven primarily by differences within the submissive group, with submissive females making significantly more chamber crossings than submissive males (p = 0.0068). No significant differences were observed between dominant males and dominant females, nor between dominant and submissive mice within each sex (p > 0.05; Fig. 6A).

**Fig.6.**
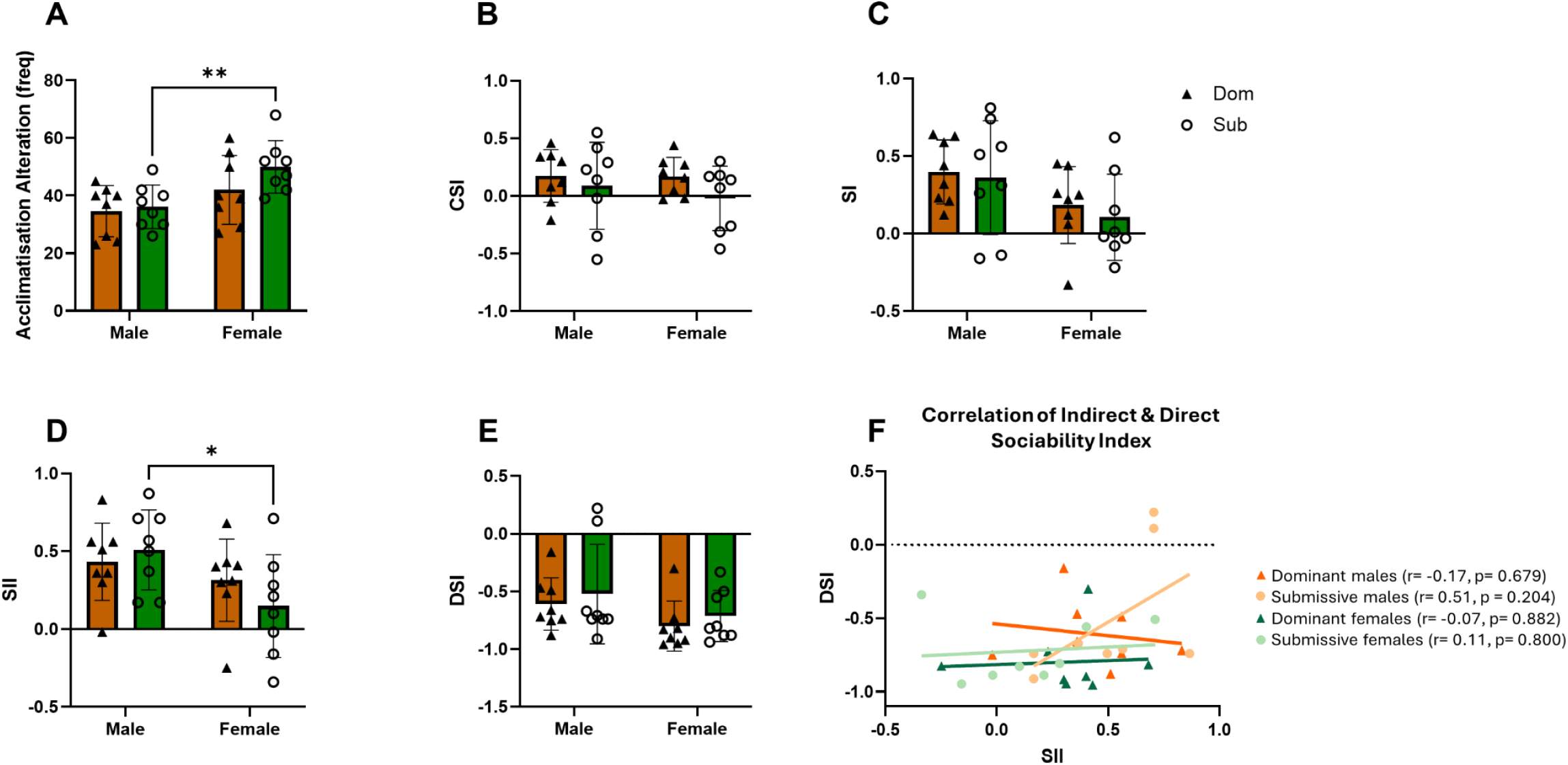
Three-chamber test results analysed based on social status in males and females. **A** Submissive females have a high number of alterations during the acclimatisation period to the arena **B** Submissive males spent more time sniffing the social cage **C** Correlation between indirect and direct social interaction across sex and social status. Scatter plots show individual data points with best-fit lines for dominant and submissive males and for dominant and submissive females. No significant correlations were found in any group (all p > 0.05) (*p<0.05, **p<0.01, two-way ANOVA, Pearson/Spearman correlation).

Two-way ANOVA revealed no significant differences across groups for the CSI (Fig. 6B), SI (Fig. 6C), or DSI (Fig. 6E). In contrast, analysis of the SII showed a significant main effect of sex (F (1, 28) = 6.010, p = 0.0207) but no main effect of social status and no sex × social status interaction was detected (p > 0.05). Post hoc comparisons showed that the sex difference was driven by a significant difference within the submissive group, with submissive males exhibiting higher SII scores than submissive females (p = 0.0144). No significant differences were found between dominant and submissive mice within either sex, and dominant males did not differ from dominant females (p > 0.05; Fig. 6D).

Correlation analyses were performed to examine the relationship between indirect and direct social interactions across social status and sex groups. No significant correlations were observed in any group (Fig. 6F).

## 4. Discussion

This study evaluated how the standard three-chamber test (3CT) captures social motivation in mice and whether adding a direct interaction phase provides a more informative measure of sociability. Guided by our systematic review of existing 3CT protocols, which revealed substantial methodological heterogeneity despite a reliance on similar cage-based outcome measures, we implemented the most commonly used protocol to ensure comparability with the literature. By modifying the task to allow physical interaction, we tested whether exploratory behaviour toward a confined conspecific reflects motivation for affiliative contact and examined how sex and social status shape these responses. Across measures, males showed greater engagement with the social stimulus, whereas females were more exploratory. These sex differences were most apparent among submissive animals. Critically, once the barrier was removed, both males and females reduced direct interaction and instead favoured exploration of the environment, suggesting the 3CT measures may overestimate affiliative social motivation. The absence of significant correlations between indirect (cage-based) and direct interaction measures further supports the conclusion that standard indices of sociability in the 3CT may overestimate sociability in mice.

### Systematic Review Reveals Methodological Variability

Our systematic review of 3CT protocols revealed substantial methodological heterogeneity across studies, despite a shared reliance on spatial and proximity-based measures to infer sociability. While most studies employed a broadly similar task structure, key parameters, including the presence and duration of acclimatisation, definitions of social proximity, and behavioural variables used to calculate sociability indices, varied considerably. However, relatively few studies explicitly accounted for baseline exploratory activity or examined how sex differences in locomotion might influence these spatial metrics. This reliance on time- and location-based measures, combined with inconsistent habituation procedures, raises concerns that variation in exploratory behaviour, rather than social motivation, may contribute substantially to reported sociability effects. These findings underscore the importance of carefully interpreting spatial indices in the 3CT and provide critical context for understanding sex-specific differences observed in the present study.

### Females showed Greater Exploration during Acclimatisation

Females exhibited more chamber crossings during acclimatisation than males, consistent with previous reports showing higher locomotor activity in female rodents under novel conditions (Augustsson et al., 2005; Chen et al., 2021). Increased exploratory behaviour in females may reflect heightened environmental scanning or sex-specific coping strategies when exposed to novelty (Knight et al., 2021; Furman et al., 2022). Importantly, this baseline difference has implications for interpreting sociability indices based on spatial measures. Greater movement between chambers may inflate or obscure measurements of social preference when time-based metrics are used alone. Thus, analyses that rely primarily on chamber occupancy may confound exploratory drive with social motivation. Exploratory behaviour and its influence on sociability measures in the 3CT may also be more sensitive to the inclusion and duration of pre-test habituation in female versus male mice.

### Standard 3CT Metrics Reflect Investigatory Interest Rather Than Direct Social Interaction

The 3CT is commonly used to assess “sociability” using indirect measures such as time spent in the chamber containing the social stimulus (Kumar & Sharma, 2016; Piccin & Contarino, 2020), proximity to the stimulus cage (Lo & Sheng, 2016; Goulding et al., 2019; Zeng et al., 2019; Nagano et al., 2018), or sniffing directed toward the cage (Juybari et al., 2020; Ferri et al., 2016). Although these variables are often interpreted as indicators of social motivation, they represent distinct behavioural processes, including spatial preference, proximity seeking, and olfactory investigation.

Consistent with several previous studies, we found no sex differences when sociability was quantified using chamber-based indices, which are particularly sensitive to baseline locomotor activity and exploratory patterns (Greene et al., 2023; Anunciado-Koza et al., 2022; Burrows et al., 2020; Lo et al., 2016). In contrast, proximity- and sniffing-based measures revealed higher cage-directed investigation in males, in line with reports that interpret increased sniffing or proximity as heightened sociability in males (Netser et al., 2017; Defensor et al., 2011; Ryan et al., 2010). However, our data extend these findings by demonstrating that such cage-directed measures primarily index investigatory interest rather than motivation for sustained social engagement.

Critically, when physical access to the stimulus mouse was permitted, both sexes reduced social interaction and increased environmental exploration. This behavioural shift indicates that earlier cage-directed behaviours reflect curiosity and information-seeking rather than affiliative social motivation. By directly comparing indirect and direct measures within the same animals, our findings challenge the assumption that increased proximity to a conspecific in a confined space equates with sociability (Beery & Shambaugh, 2021) and support the interpretation that the standard 3CT predominantly measures social curiosity rather than true social interaction.

### Latency and Frequency Measures Do Not Explain Sex Differences in Social Indices

Importantly, the observed sex differences in social indices were not driven by differences in basic approach behaviour. Males and females did not differ in latency to enter the social chamber, latency to approach the social stimulus during either the indirect or direct phases, nor in the frequency of social chamber entries, cage visits, cage-directed sniffing events, or direct interaction bouts. These null effects indicate that both sexes were equally likely to detect, approach, and initiate contact with the social stimulus. Thus, sex differences observed in proximity- and sniffing-based indices cannot be attributed to differences in social approach speed or opportunity, but instead reflect qualitative differences in how social and environmental priorities are allocated once the stimulus is encountered.

This dissociation is particularly informative because latency and frequency measures are often interpreted as indicators of motivation or salience. Our findings suggest that such measures may be insensitive to meaningful differences in social strategy, especially when animals readily approach the stimulus but differ in how long they sustain investigatory or interactive behaviours. Together, these results reinforce the conclusion that commonly used 3CT metrics do not reflect a unitary construct of sociability and that differences in social indices arise from behavioural prioritisation rather than from deficits or enhancements in basic social approach.

### Cage-Directed Investigation does not Predict Physical Social Interaction during Direct Contact

The addition of a direct contact phase provided behavioural information that is not accessible using the standard three-chamber test. During unrestricted access, males continued to engage more than females in both social investigation and direct contact behaviours, consistent with reports of higher male social engagement (Ma et al., 2022) or dominance-related motivation (Borak et al., 2025). However, despite this sex difference, both males and females redirected their behaviour away from the social partner toward environmental exploration once the barrier was removed, indicating that exploratory drive remains a dominant behavioural priority even in the presence of a social opportunity (Kikuchi et al., 2022).

Together, these findings demonstrate that investigatory interest toward a confined conspecific does not necessarily translate into sustained social interaction when physical access is permitted. This pattern contrasts with observations in rats, where direct social interaction paradigms typically elicit prolonged engagement in social behaviours (Varlinskaya & Spear, 2008), highlighting potential species differences and the importance of assay design.

Importantly, indirect and direct measures were not correlated across sex or dominance groups, indicating that cage-directed investigation and physical interaction reflect distinct behavioural dimensions rather than a single underlying construct. Indirect metrics appear to index curiosity-driven social investigation, whereas direct interaction more accurately reflects affiliative engagement. Integrating both approaches, therefore, provides a more robust and biologically meaningful framework for characterising social behaviour in mice.

### Sex Differences are Expressed in Submissive Mice

Sex differences were primarily expressed among submissive mice. Submissive females made more chamber crossings during acclimatisation, suggesting greater locomotor reactivity to novelty, whereas submissive males displayed lower exploration. These behavioural profiles likely reflect sex-specific coping strategies under social subordination (Williamson et al., 2019; Frischknecht et al., 1982). In contrast, submissive males exhibited higher social interest scores than submissive females, driven by increased sniffing behaviour. Rather than reflecting enhanced sociability, this pattern may reflect a risk-minimising strategy for social assessment (Matzel et al., 2017). Male mice frequently use olfactory cues to monitor competitors and assess hierarchical status (Kikusui, 2013; Karlsson et al., 2015), and increased sniffing may therefore reflect vigilance rather than affiliation.

Although social status did not produce a main effect in the overall analysis, post hoc tests revealed that sex differences emerged only among subordinate animals. Dominant mice showed no sex divergence, suggesting that hierarchy stabilises social strategies (So et al., 2015), whereas submissive animals exhibit more variable and sex-dependent behaviours (Watanabe, 2014; Fulenwider et al., 2024). Importantly, relying on sniffing alone to infer sociability risks misclassification. Elevated sniffing by submissive males could be misinterpreted as prosocial behaviour when it may instead reflect information gathering under perceived threat (Wesson, 2013). The absence of overt aggression supports the interpretation that behaviour was investigative rather than competitive (van Loo et al., 2004). Females, by contrast, showed stronger environmental prioritisation, consistent with reports that social engagement is less dominant in female behavioural repertoires under laboratory conditions (Granza et al., 2023).

### Implications and Limitations

Our findings suggest that the 3CT, while valuable for measuring social approach, does not capture the full complexity of social behaviour in mice. Chamber time is influenced by locomotor activity, cage proximity reflects investigatory interest, and only direct interaction measures provide insight into affiliative behaviour (Lecker et al., 2021). The modified 3CT demonstrates that exploratory motivation predominates once physical access is allowed, challenging the assumption that interest in a confined conspecific equates to social motivation.

Several methodological limitations should be acknowledged. The direct contact phase was brief, potentially limiting observation of stable interaction patterns. Extending this period may reveal stronger effects related to affiliation or dominance. Additionally, the study did not include physiological or neural markers that could inform motivation or arousal. Integrating measures such as ultrasonic vocalisations (Premoli et al., 2022), stress and sex hormones (Chari et al., 2020; Nisbett et al., 2023) may further clarify behavioural interpretations. Finally, testing dominance-matched or mismatched dyads would provide further insight into hierarchical modulation of social behaviour (Varholick et al., 2019).

## Conclusion

In conclusion, our results show that the standard three-chamber test primarily measures social investigation rather than affiliative behaviour and is strongly influenced by exploratory activity. Sex differences in social indices are driven largely by submissive animals and do not generalise across dominance status. The inclusion of a direct interaction phase reveals a dissociation between investigatory interest and social engagement, highlighting the need for behavioural assays that combine indirect measures with direct, unrestricted social interaction to more accurately characterise sociability in mice. Together, these findings add to a growing body of literature challenging the assumption that mice consistently seek social contact driven by affiliative motivation and underscore the importance of re-evaluating how social behaviour is operationally defined and measured.

## Acknowledgements

The authors thank the staff of the Animal Services Unit at the University of Bristol for their support with animal care and facility management. We also thank our laboratory colleagues for their technical assistance and helpful discussions throughout the study.

## CRediT Author Contribution Statement

**MS:** Conceptualisation, Methodology, Investigation, Formal analysis, Data Curation, Visualisation, Writing – Original Draft. **HDP:** Supervision, Project administration, Writing – Review & Editing, Funding Acquisition. **MGJ:** Conceptualisation, Methodology, Software, Supervision, Resources, Writing – Review & Editing. **ESJR:** Conceptualisation, Methodology, Supervision, Project administration, Data curation, Writing – Review & Editing.

## Funding

This work was supported by the Indonesia Endowment Fund for Education and by institutional support from the University of Bristol.

## Conflict of Interest Statement

ESJR and MGJ are creators of the 3Hs initiative, a framework designed to promote refined housing, handling and habituation methods. This initiative includes working with commercial suppliers of products used in the management of laboratory animals. ESJR has received funding for collaborative and contract research from pharmaceutical companies, including Boehringer Ingelheim, Compass Pathways, Eli Lilly, IRLabs Therapeutics, Pfizer, and SmallPharma, and has acted as a paid consultant for Compass Pathways and Pangea Botanicals.

## Notes

### Competing Interest Statement

The authors have declared no competing interest.

