## Supplementary Information for "Beyond the Barrier: An Assessment of Social Behaviour in Mice using a Modified 3-Chamber Test"

**Figure 1**

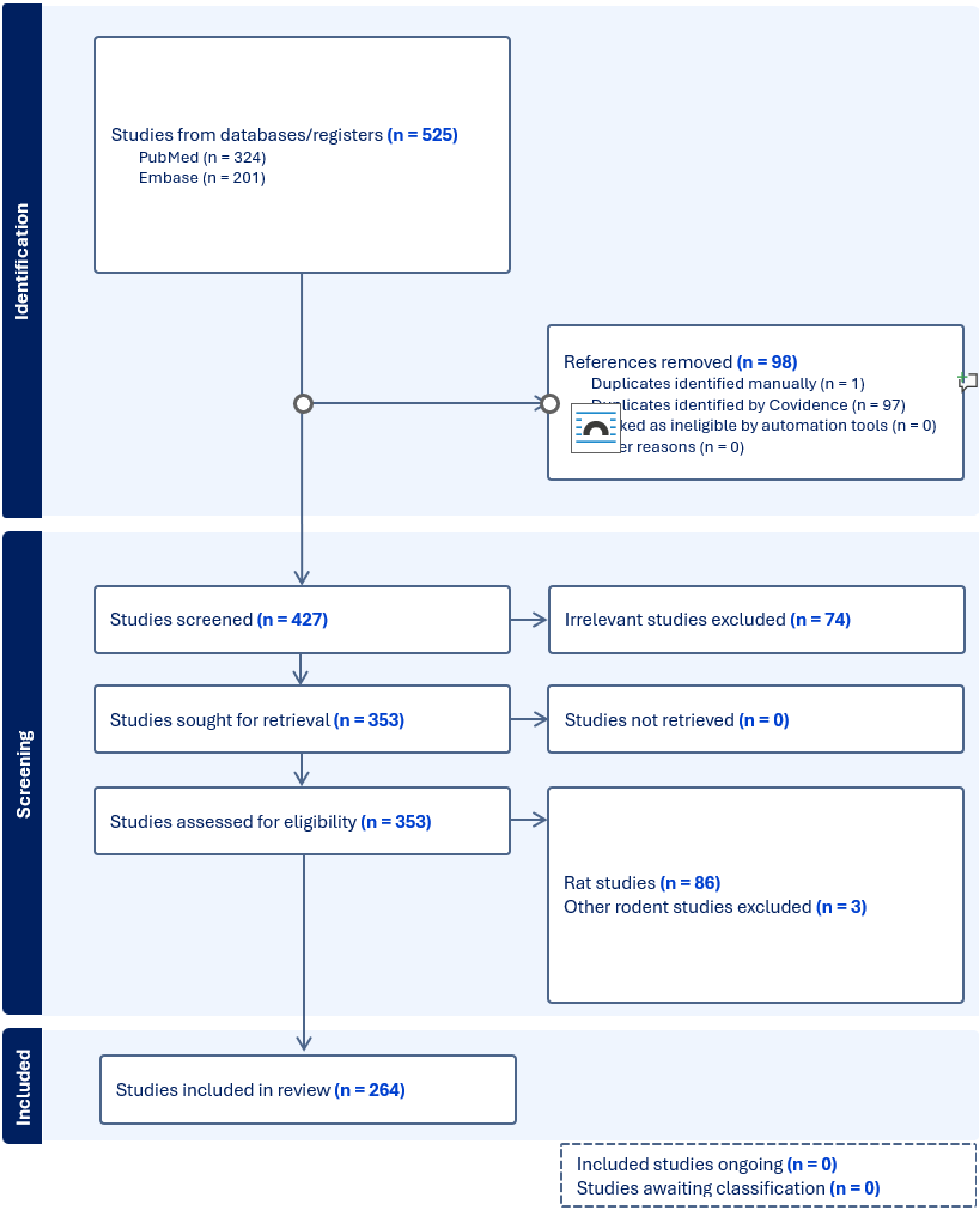


***Fig.1.*** **PRISMA data**. Flowchart of the study selection process for systematic review.

**Figure 2**

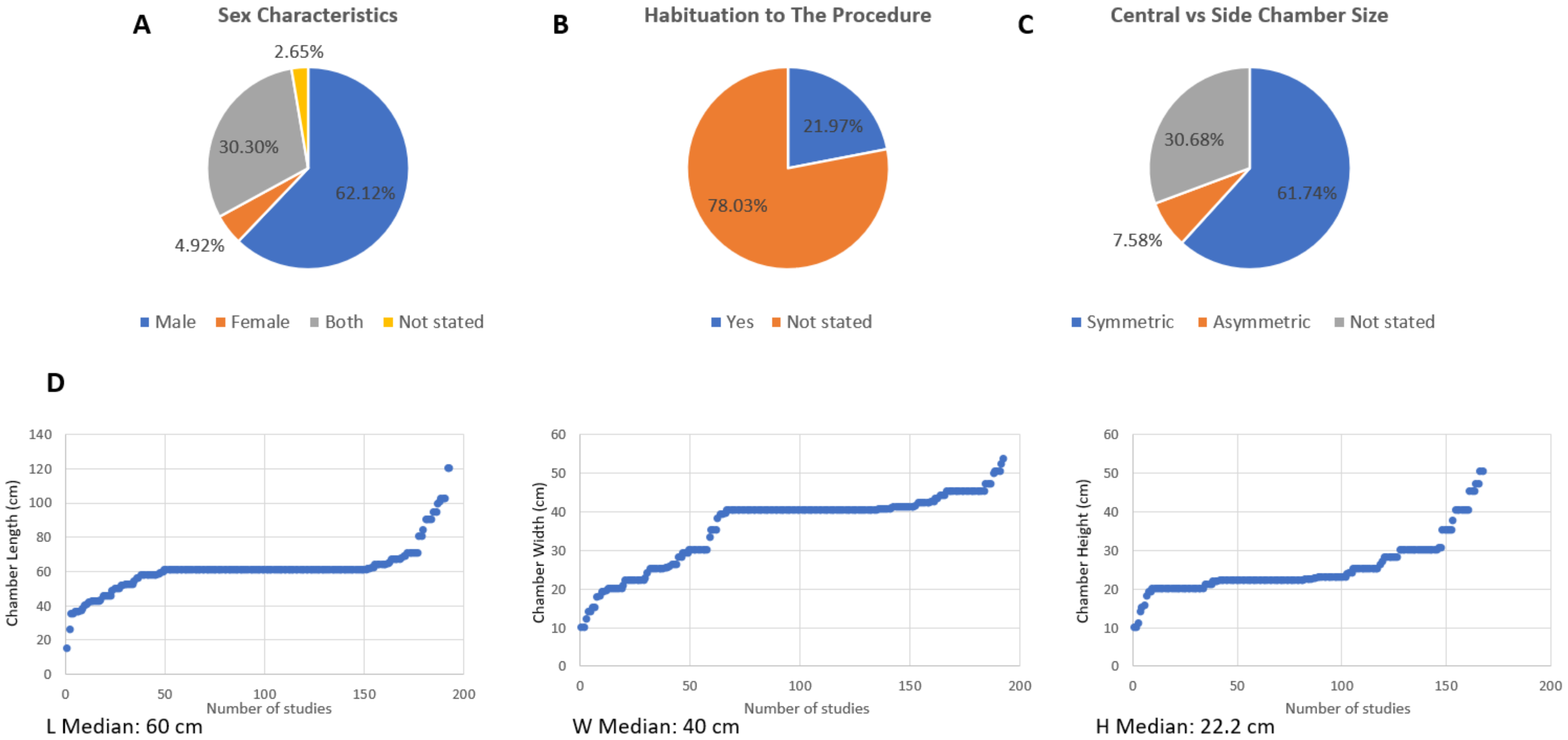


***Fig.2. The data obtained from studies using the three-chamber test in mice. A** Most sociability studies use male-only. **B** Most studies did not habituate the mouse to the arena and the cage. **C** Most studies use equal chamber size between the central and side chambers. **D** Variation in the three-chamber apparatus size between studi*

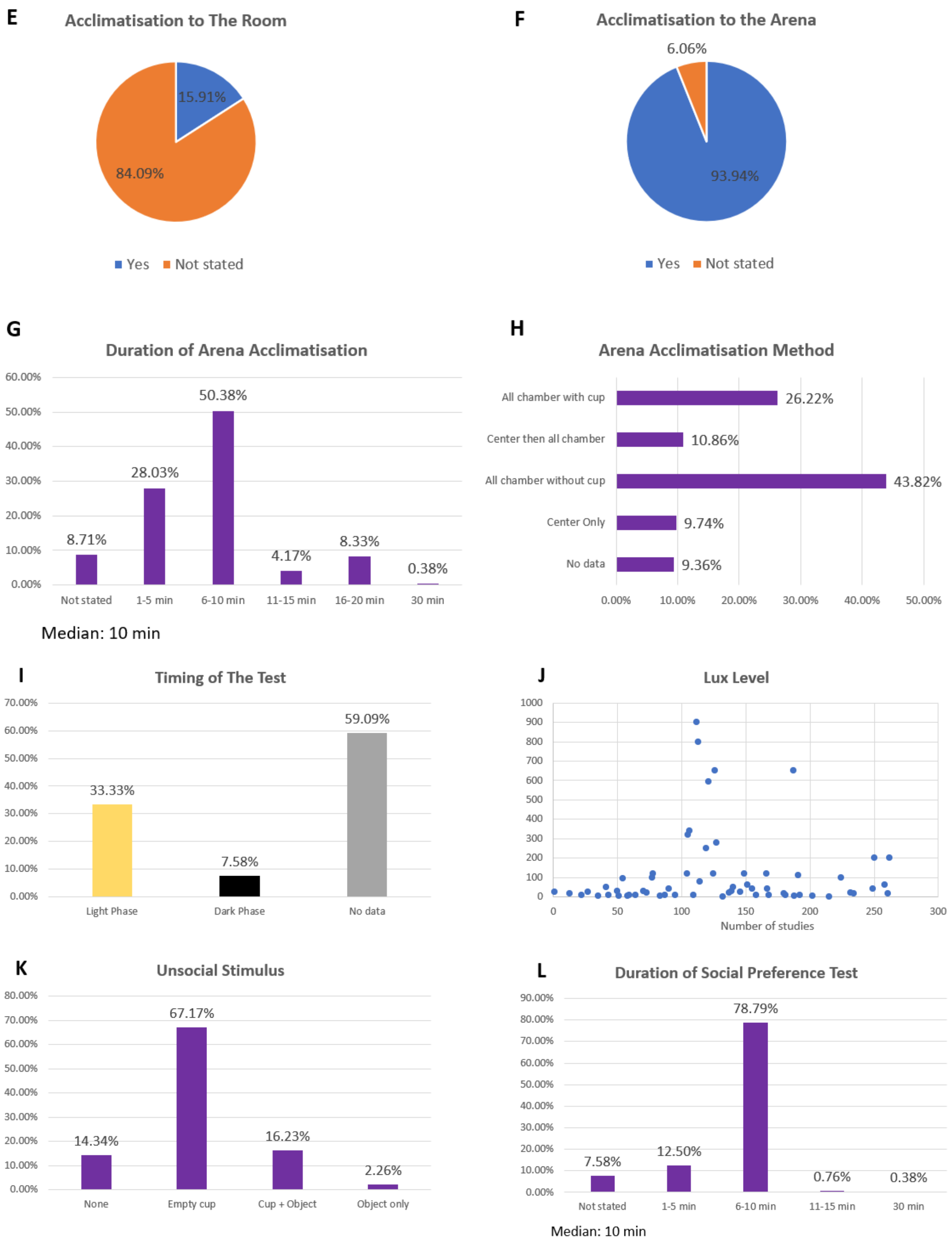


***Fig.2. The data obtained from studies using the three-chamber test in mice (cont). E*** *Most studies did not habituate or acclimatise the mouse to the behaviour room before the test, but* ***F*** *acclimatised the test mouse to the arena.* ***G*** *Acclimatisation was done for 6-10 minutes.* ***H*** *The most common acclimatisation method was allowing the test mouse to explore all empty chambers.* ***I*** *Most studies did not state the timing of the test* ***J*** *Variation in the brightness of the behaviour room, ranging from 2 to 900 lux.* ***K*** *The most common method for the social preference test was using the empty cage as the unsocial stimulus.* ***L*** *The duration of the social preference test was 6-10 minutes.*

**Figure 3**

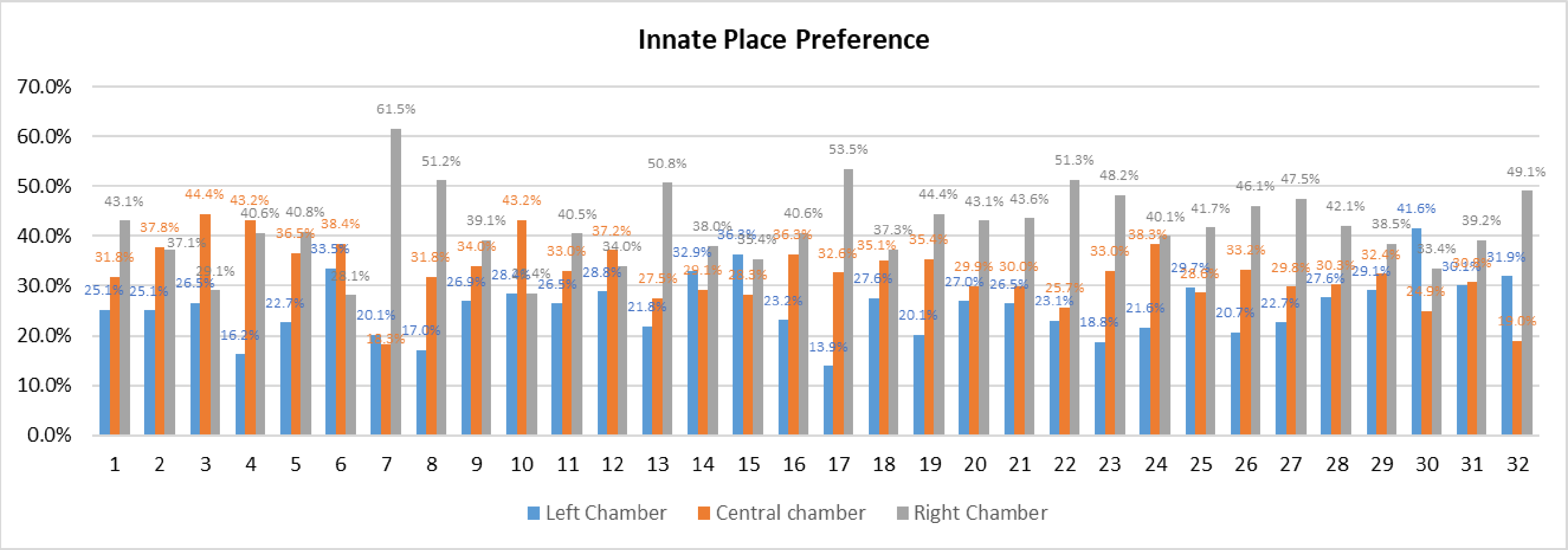


***Fig.3.*** *None of the test mice have an innate place preference (IPP) during acclimatisation, as they spent <80% of total acclimatisation time in all three chamber*

**Figure 4**

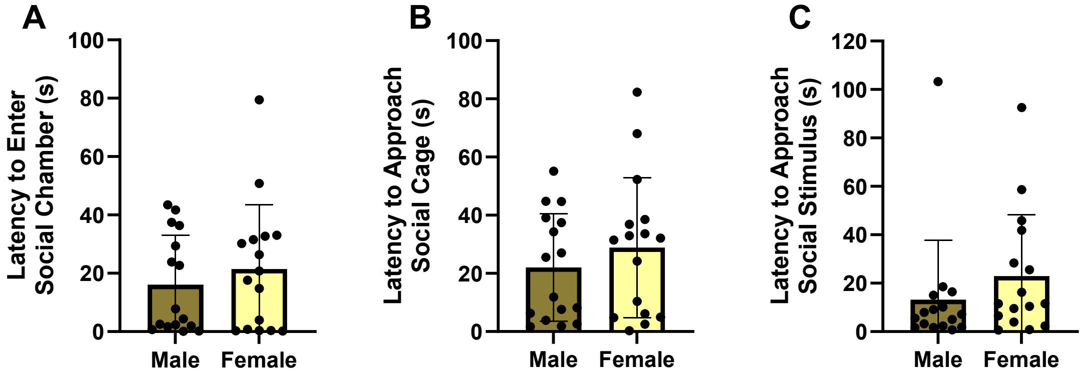


***Fig. 4.*** *There was no difference between sex groups in their* ***A*** *latency to enter the social chamber,* ***B*** *in latency to approach the social cage, and* ***C*** *in latency to approach social stimulus (p>0.05, independent t-test).*

**Figure 5**

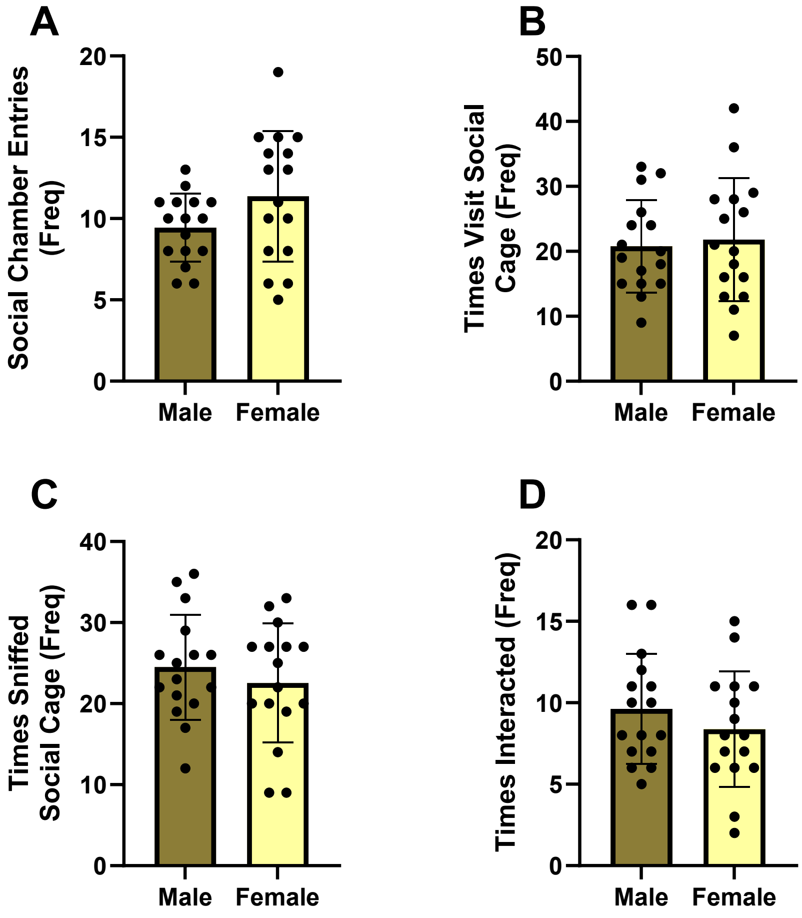


***Fig.5.*** *Between males and females, there was no difference in* ***A*** *frequency entering the chamber containing the stimulus mouse,* ***B*** *frequency visiting the cage containing the stimulus mouse,* ***C*** *frequency sniffing the social cup, and* ***D*** *frequency interacting after the cage was lifted (p>0.05, independent t-test).*
